## Supplementary information for "Harnessing the regenerative potential of *interleukin11* to enhance heart repair"

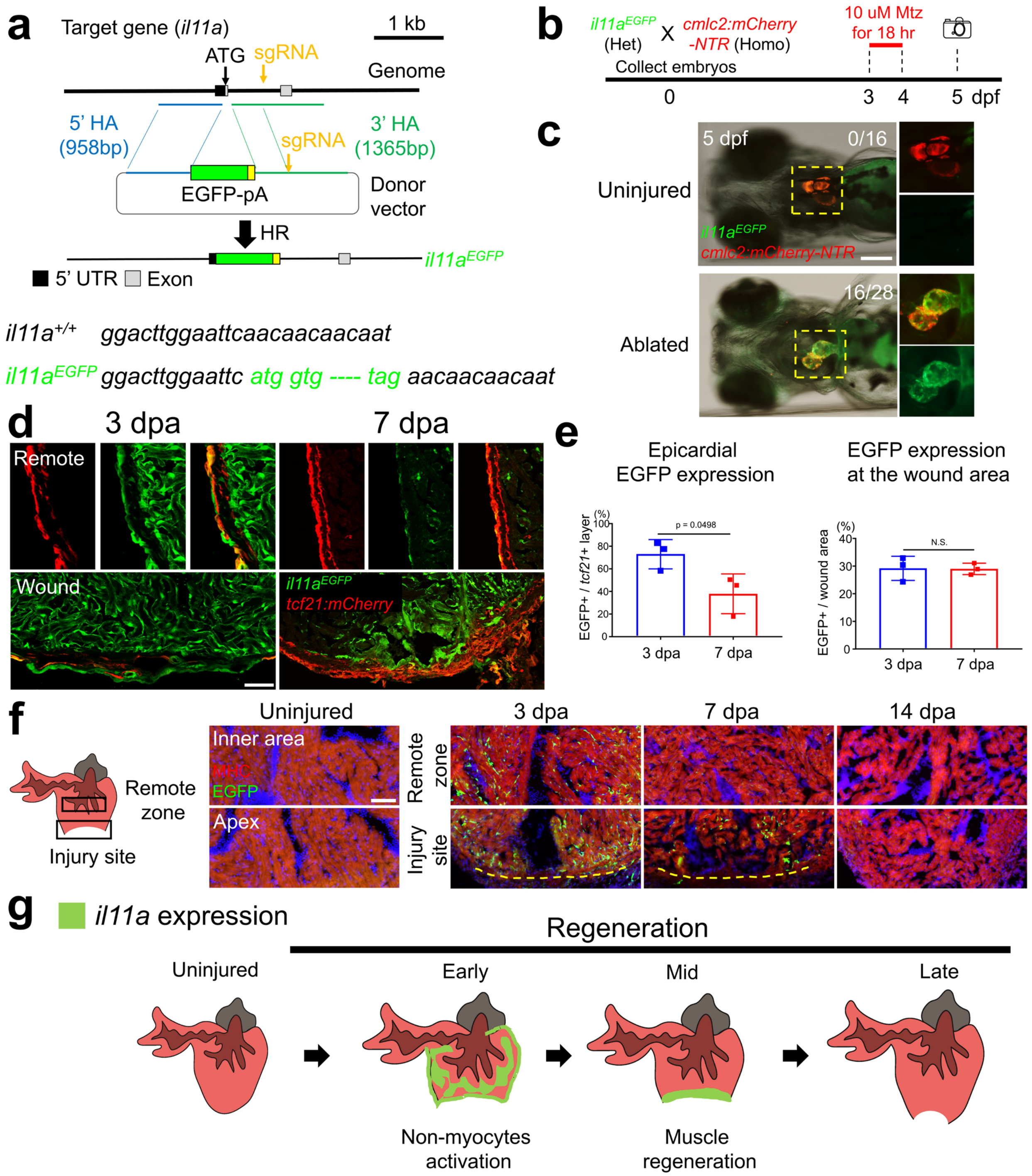


**Supplementary Fig. 1. *il11a* is not expressed at the early development, but during regeneration in zebrafish heart.**

(**a**) Genomic structure of the endogenous *il11a* gene and the reporter gene. sgRNA and the *EGFP* reporter gene together with Cas9 protein were injected into embryos, and a transgenic line containing the *EGFP* gene in the *il11a* locus was established. ATG, Transcription start site, HA, Homologous Arms, HR, Homologous Recombination. (**b**) Zebrafish strains and experimental design employed to examine *il11a*$^{EGFP}$ expression in uninjured and injured hearts of larvae. Het, Heterozygote. Homo, Homozygote. (**c**)

Representative images of uninjured and ablated hearts in *il11a*$^{EGFP}$ larvae at 5 days-post fertilization (dpf). Insets correspond to higher magnifications of dashed boxes. The number in the upper right corner of each image represents the fraction of the fish expressing EGFP. (**d**) Representative cardiac section images of *il11a*$^{EGFP}$ and *tcf21:mCherry* in 3 dpa (left) and 7 dpa (right) hearts. (**e**) Quantification of *EGFP* expression area in *tcf21*$^{+}$ epicardium (left) and at the wound area (right) at 3 dpa and 7 dpa. n = 3. (**f**) Representative images of *EGFP* expression at the injury site and remote zone in *i11a*$^{EGFP}$ hearts during heart regeneration. MHC (red) indicates myocardium. (**g**) Spatiotemporal expression and potential roles of *il11a*. *il11a* is strongly induced throughout the ventricle at the early stage of regeneration to activate non-cardiac muscle cells. At the intermediate stage, *il11a* expression is restricted to the wound area to promote CM proliferation. *il11a* expression returns to the basal level at the late stage of the regeneration. Scale bar, 100 μm in **c** and 50 μm in **d** and **f**.

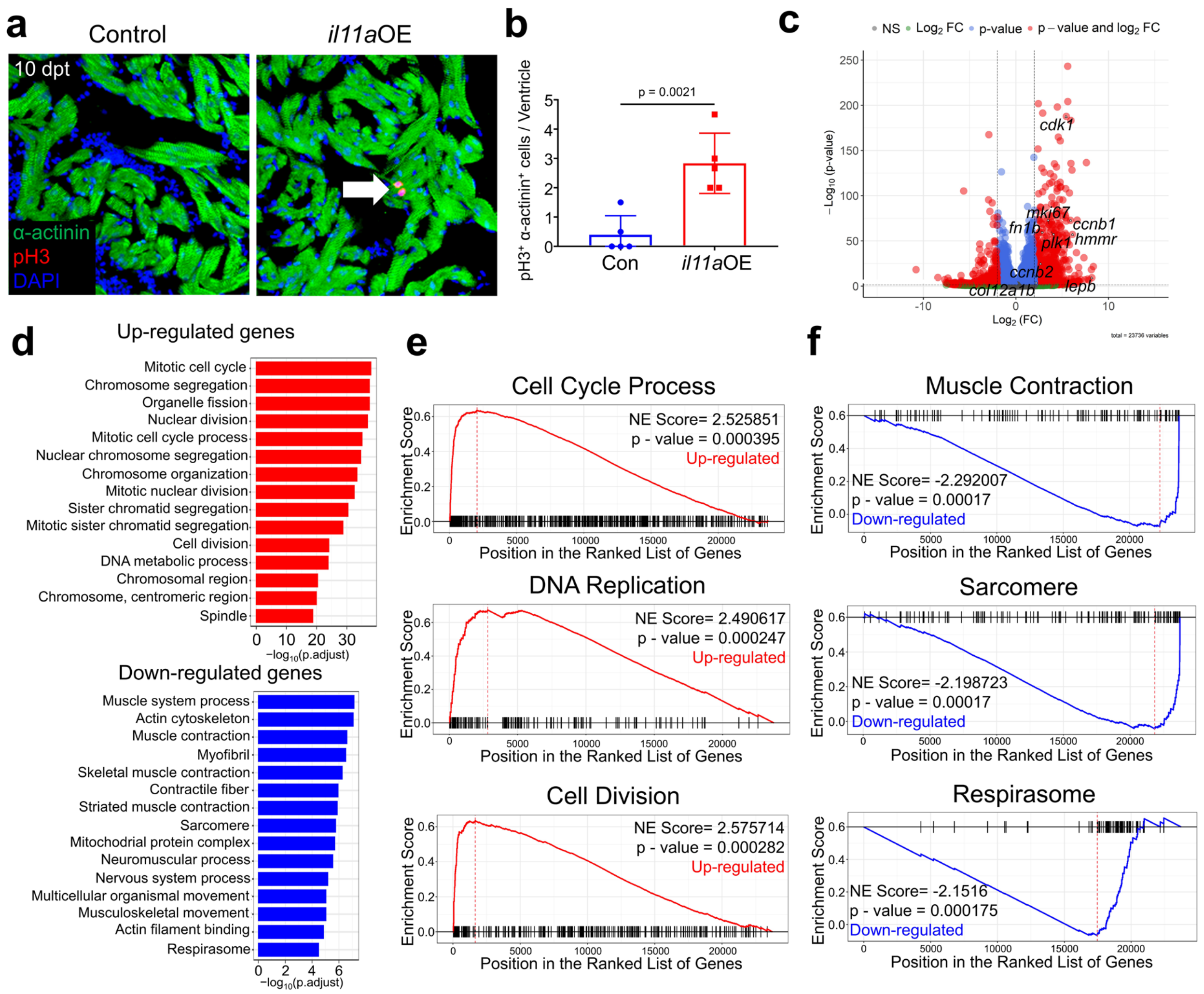


**Supplementary Fig. 2. *il11a*OE triggers CM dedifferentiation and proliferation at transcriptomic level without injury signal.**

(**a**) Representative image of cardiac sections stained with α-actinin (green, CM) and phosphorylated-histone3 (pH3, red) from control and *il11a*OE uninjured hearts at 10 dpt. An arrow indicates pH3⁺ CMs. (**b**) Quantification of pH3⁺ CMs in the whole ventricle. n = 5. (**c**) Differentially expressed genes between control and *il11a*OE ventricles shown as a volcano plot. (**d**) Gene ontology (GO) enrichment analysis of up- (top) and down- (bottom) regulated genes in the *il11a*OE. The bars indicate the adjusted P-value for the gene enrichment in our analysis. (**e, f**) Gene Set Enrichment Analysis (GSEA) plots of the up-regulated (**e**) and down-regulated (**f**) genes from control and *il11a*OE. *il11a*OE upregulates gene expression associated with cell cycle activity, DNA replication, and cell division while downregulated genes are associated with cardiac muscle contraction, sarcomere, and respirasome. Scale bar, 50 μm in **a**.

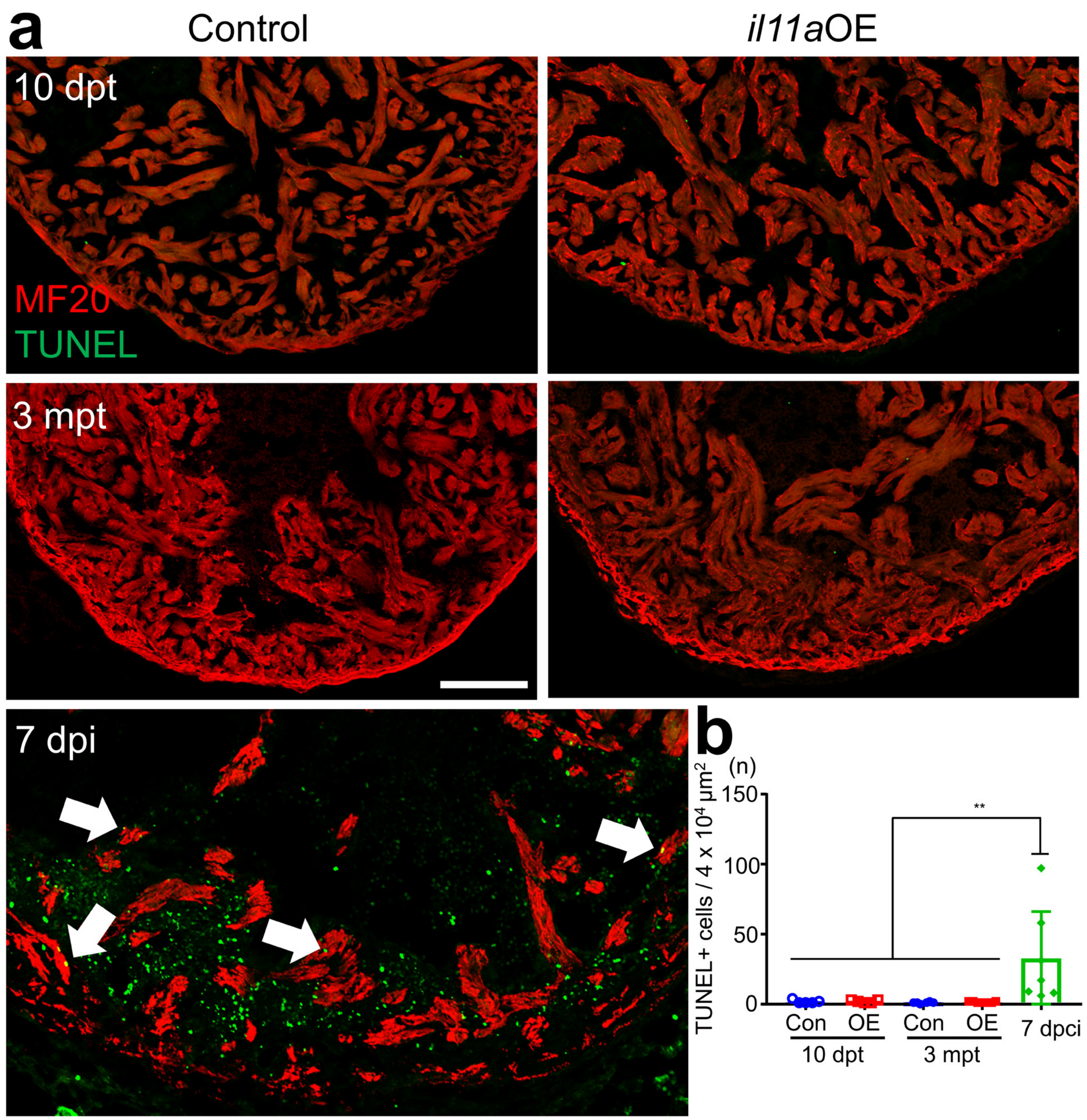


**Supplementary Fig. 3. *il11a*OE does not induce apoptosis and cell death in CMs.** (**a**) Representative cardiac section images stained with MF20 (Red) from Con and *il11a*OE at 10 days post-treatment (dpt) and 3 months post-treatment (mpt). Apoptotic cells are detected by TUNEL assay (Green). 7 days post-cryoinjury (dpi) hearts are used as a positive control. Arrows indicate apoptotic CMs. (**b**) Quantification of the number of apoptotic cells. n = 6 - 7. Scale bars, 100 μm in **a**.

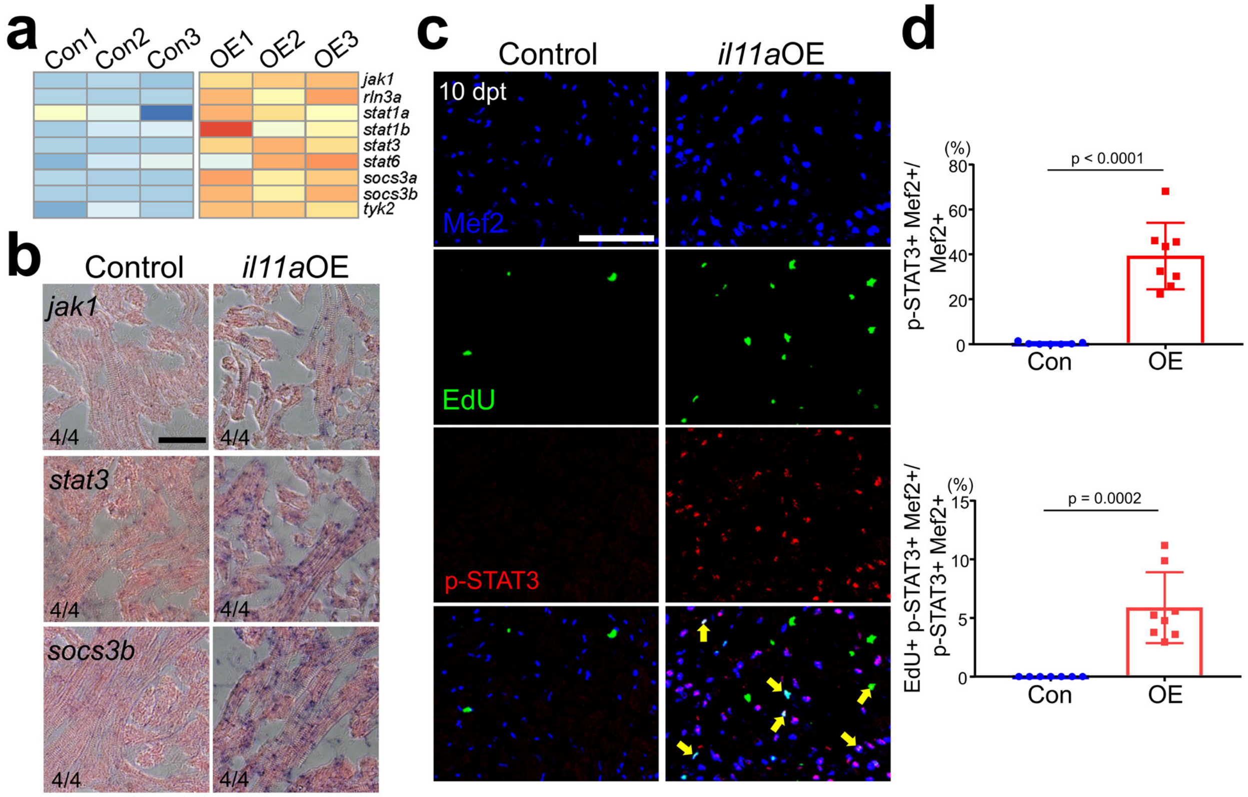


**Supplementary Fig. 4. *il11a*OE activates JAK/STAT pathway in CMs to induce proliferation.**

(**a**) Heatmap of differential gene expression associated with JAK/STAT pathway for control and *il11a*OE. (**b**) Representative images of *in situ* hybridization (ISH) on cardiac sections of control and *il11a*OE. The number in the lower left corner of each image represents the fraction of the analyzed hearts with displayed phenotype. n = 4. (**c**) Representative images of cardiac sections stained with Mef2 (Blue), EdU (Green), and p-STAT3 (red) from control and *il11a*OE uninjured hearts at 10 dpt. Arrows indicate p-STAT3$^+$ EdU$^+$ CMs. (**d**) (Top) The quantification graph of p-STAT3$^+$ Mef2$^+$ CMs out of all Mef2$^+$ CMs. (Bottom) The quantification graph of EdU$^+$ pSTAT-3$^+$ Mef2$^+$ CMs out of pSTAT-3$^+$ Mef2$^+$ CMs. n = 7 - 8. Scale bar, 20 μm in **b** and 50 μm in **c**.

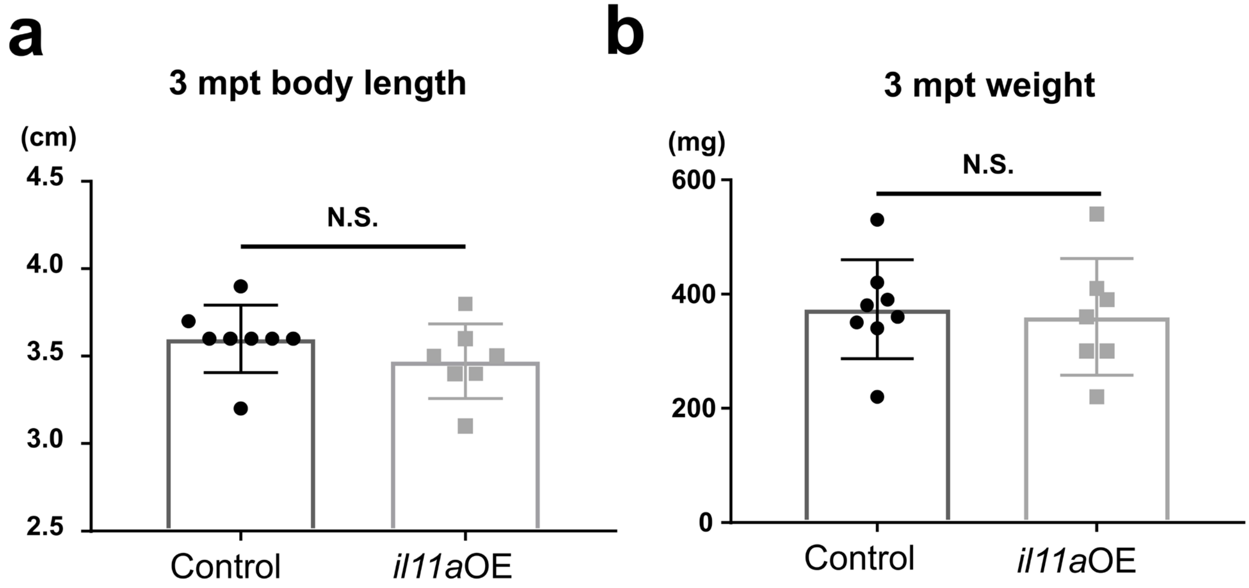


**Supplementary Fig. 5. *il11a*OE adult fish have similar body length and weight to control.**
(**a, b**) The measurement of adult zebrafish body length (**a**) and body weight (**b**) 3 months after 4-HT treatment yielded no significant difference between control and *il11a*OE. n = 7 - 8.

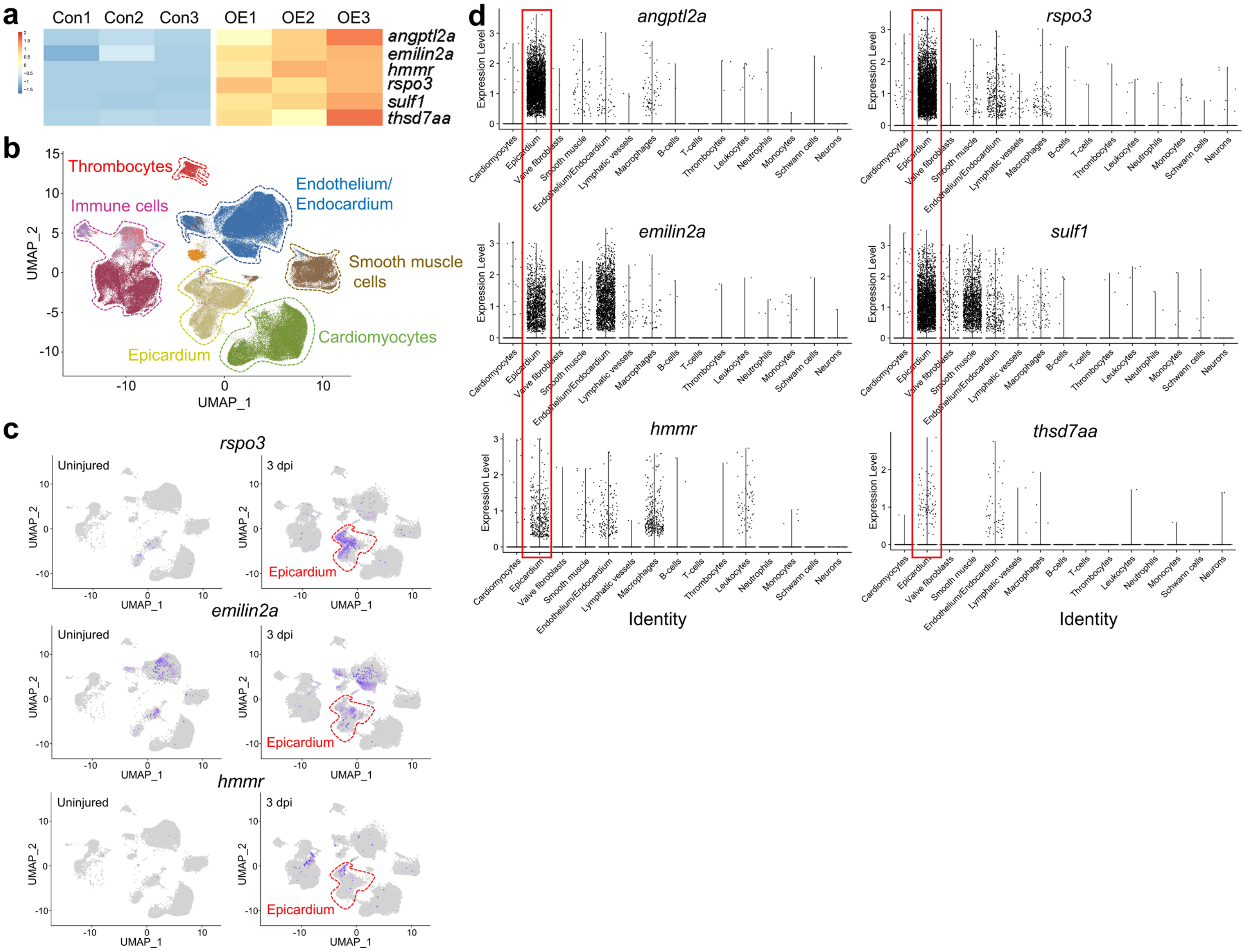
a
Con1 Con2 Con3 OE1 OE2 OE3
angptl2a
emilin2a
hmmr
rspo3
sulf1
thsd7aa
b
Thrombocytes
Immune cells
Endothelium/
Endocardium
Smooth muscle
cells
Epicardium
Cardiomyocytes
UMAP_1
UMAP_2
c
rspo3
emilin2a
hmmr
Uninjured
3 dpi
Epicardium
d
angptl2a
rspo3
emilin2a
sulf1
hmmr
thsd7aa
Expression Level
Cardiomyocytes
Epicardium
Valve fibroblasts
Smooth muscle
Endothelium/Endocardium
Lymphatic vessels
Macrophages
B-cells
T-cells
Thrombocytes
Leukocytes
Neutrophils
Monocytes
Schwann cells
Neurons
Identity

**Supplementary Fig. 6. *il11a*OE stimulates coronary growth by EPC-mediated angiogenic factors.**
(**a**) Heatmap of differential gene expression responsible for vascularization from control and *il11a*OE. (**b**) Uniform manifold approximation and projection (UMAP) plot of zebrafish regenerating hearts and cell cluster analysis. (**c**) Gene expression plots of *rspo3, emilin2a*, and *hmmr* induced in epicardium at 3 days post injury (dpi). Cells expressing indicated genes are colored purple, and the relative intensity indicates relative expression levels. Arrows indicate epicardium-specific expression of indicated genes. (**d**) Violin plot showing expression of angiogenic factors in the cell populations of the hearts. Red boxes indicate the expression of the indicated genes in epicardium.

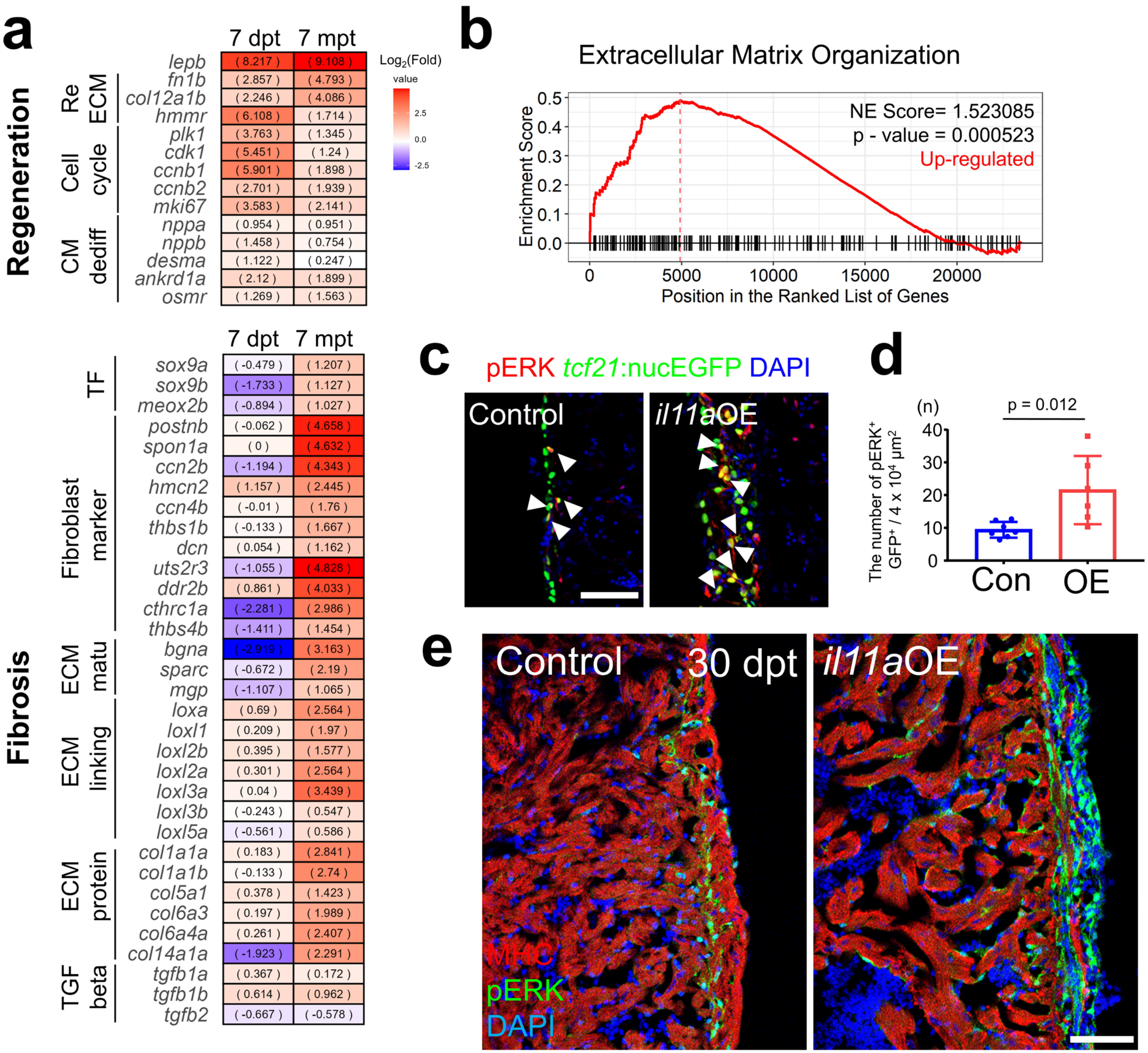
a
Regeneration
Fibrosis
7 dpt 7 mpt
Log2(Fold)
value
2.5
0.0
-2.5
Re ECM
Cell cycle
CM dediff
lepb (8.217) (9.108)
fn1b (2.857) (4.793)
col12a1b (2.246) (4.086)
hmmr (6.108) (1.714)
plk1 (3.763) (1.345)
cdk1 (5.451) (1.24)
ccnb1 (5.901) (1.898)
ccnb2 (2.701) (1.939)
mki67 (3.583) (2.141)
nppa (0.954) (0.951)
nppb (1.458) (0.754)
desma (1.122) (0.247)
ankrd1a (2.12) (1.899)
osmr (1.269) (1.563)
TF
Fibroblast marker
ECM matu
ECM linking
ECM protein
TGF beta
sox9a (-0.479) (1.207)
sox9b (-1.733) (1.127)
meox2b (-0.894) (1.027)
postnb (-0.062) (4.658)
spon1a (0) (4.632)
ccn2b (-1.194) (4.343)
hmcn2 (1.157) (2.445)
ccn4b (-0.01) (1.76)
thbs1b (-0.133) (1.667)
dcn (0.054) (1.162)
uts2r3 (-1.055) (4.828)
ddr2b (0.861) (4.033)
cthrc1a (-2.281) (2.986)
thbs4b (-1.411) (1.454)
bgna (-2.919) (3.163)
sparc (-0.672) (2.19)
mgp (-1.107) (1.065)
loxa (0.69) (2.564)
loxl1 (0.209) (1.97)
loxl2b (0.395) (1.577)
loxl2a (0.301) (2.564)
loxl3a (0.04) (3.439)
loxl3b (-0.243) (0.547)
loxl5a (-0.561) (0.586)
col1a1a (0.183) (2.841)
col1a1b (-0.133) (2.74)
col5a1 (0.378) (1.423)
col6a3 (0.197) (1.989)
col6a4a (0.261) (2.407)
col14a1a (-1.923) (2.291)
tgfb1a (0.367) (0.172)
tgfb1b (0.614) (0.962)
tgfb2 (-0.667) (-0.578)
b
Extracellular Matrix Organization
Enrichment Score
NE Score= 1.523085
p - value = 0.000523
Up-regulated
0.5
0.4
0.3
0.2
0.1
0.0
0
5000
10000
15000
20000
Position in the Ranked List of Genes
c
pERK tcf21:nucEGFP DAPI
Control
il11aOE
d
(n)
p = 0.012
The number of pERK+ GFP+ / 4 x 10^4 μm^2
40
30
20
10
0
Con
OE
e
Control
30 dpt
il11aOE
MHC
pERK
DAPI

**Supplementary Figure 7. pERK-mediated epicardial activation ultimately results in fibrosis in *il11a*OE hearts.**

(**a**) Differential expression of genes associated with regeneration (top) and cardiac fibrosis (bottom) between 7 dpt and 7 mpt. Genes associated with cardiac fibroblasts and pathological fibrosis are significantly upregulated in 7 mpt, but not 7 dpt, *il11a*OE, compared to control. In contrast to that, pro-regenerative genes, such as cell cycle and regenerative ECM, are highly upregulated in both 7 dpt and 7 mpt *il11a*OE. Re-ECM, regenerative ECM; CM dediff., CM dedifferentiation; TF, transcription factor; ECM matu., ECM maturation. (**b**) GSEA plots of the extracellular matrix organization from control and *il11a*OE. At 7 mpt, ECM organization-related genes are noticeably upregulated in *il11a*OE, compared to control. (**c**) Representative cardiac section images of 30 dpt Con and *il11a*OE hearts. Green and red indicate *tcf21*:nucEGFP and pERK, respectively. Arrowheads represent $pERK^+$ *tcf21:*$nucEGFP^+$ epicardial cells. (**d**) The number of $pERK^+$ *tcf21*:$nucGFP^+$ epicardial cells between Con and *il11a*OE. n = 7 - 8 (**e**) Representative image of heart sections stained with CM (MHC, red) and pERK (green) from Con and *il11a*OE at 30 dpt. n = 3. pERK is significantly induced in ventricular wall, but not in $MHC^+$ myocardium by *il11a*OE. Scale bar, 50 μm in **c** and **e**.

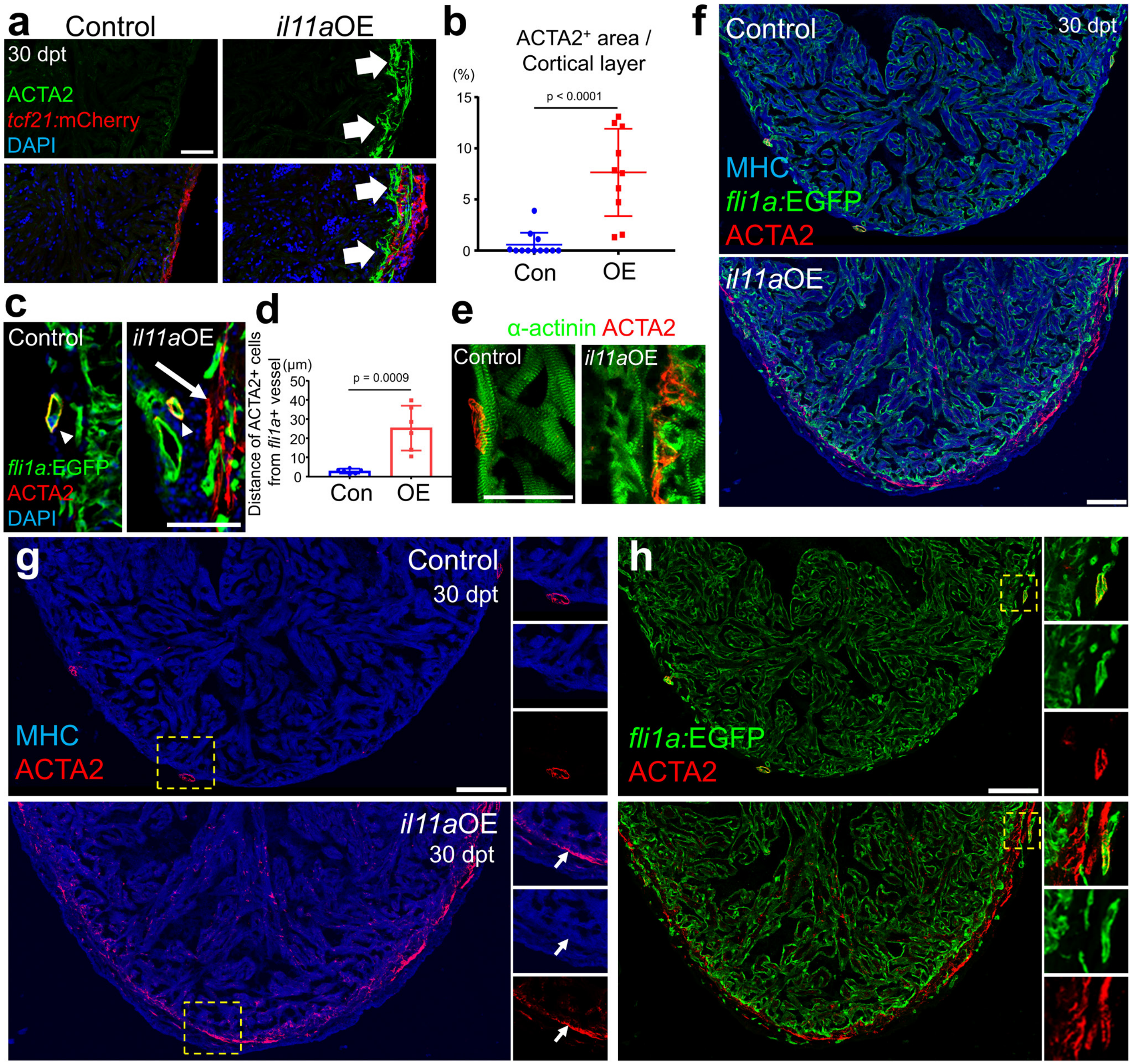
a
Control
il11aOE
30 dpt
ACTA2
tcf21:mCherry
DAPI
b
ACTA2+ area / Cortical layer
(%)
15
10
5
0
p < 0.0001
Con
OE
c
Control
il11aOE
fli1a:EGFP
ACTA2
DAPI
d
Distance of ACTA2+ cells from fli1a+ vessel
(μm)
50
40
30
20
10
0
p = 0.0009
Con
OE
e
α-actinin ACTA2
Control
il11aOE
f
Control
30 dpt
MHC
fli1a:EGFP
ACTA2
il11aOE
g
Control
30 dpt
MHC
ACTA2
il11aOE
30 dpt
h
fli1a:EGFP
ACTA2

**Supplementary Figure 8. ACTA2 is expressed in vascular smooth muscle cells and dedifferentiating cardiomyocytes.**

(**a**) Representative cardiac section images of 30 dpt control and *il11a*OE hearts. Green and red indicate ACTA2$^+$ cell layer and *tcf21*$^+$ epicardium, respectively. Arrows represent ACTA2$^+$ area. (**b**) Quantification of ACTA2$^+$ area in the cortical layer of the uninjured hearts between Con and *il11a*OE at 30 dpt. n = 10 - 12. (**c**) Representative cardiac section images of 30 dpt Con and *il11a*OE hearts. Green and red indicate *fli1a*:EGFP and ACTA2, respectively. *il11a*-induced ACTA2+ cells are localized distantly to *fli1a*:EGFP+ area in *il11a*OE. Encircling ACTA2+ in control and il11a OE hearts display close proximity to *flia1*:EGFP+ endothelial cells in a coronary vessel, indicating VSMCs. (**d**) Quantification of distance between ACTA2+ cells and *flia1*:EGFP+ area for Con and *il11a*OE. n = 6 (**e**) Representative cardiac section images of 30 dpt control and *il11a*OE hearts stained with α-actinin (CM, green) and ACTA2 (red), respectively. n = 6. *il11a*OE shows distinct ACTA2$^+$ cell populations from α-actinin$^+$ CMs although partial overlapping of ACTA2 signal with α-actinin$^+$ CMs are observed. (**f**) Cardiac section images of 30 dpt control and *il11a*OE hearts stained with *fli1a*:EGFP (green), ACTA2 (red), and MHC (blue), respectively. **(g)** and **(h)** show double channel images for green ACTA2 (red); MHC (blue) and *fli1a*:EGFP (green); ACTA2 (red), respectively. Insets on the right side correspond to higher magnifications of dashed boxes. Arrows indicate colocalization of MHC$^+$ CM and ACTA2$^+$ cells. Scale bar, 50 μm in **a, c,** and **e** and 100 μm in **f, g,** and **h**.

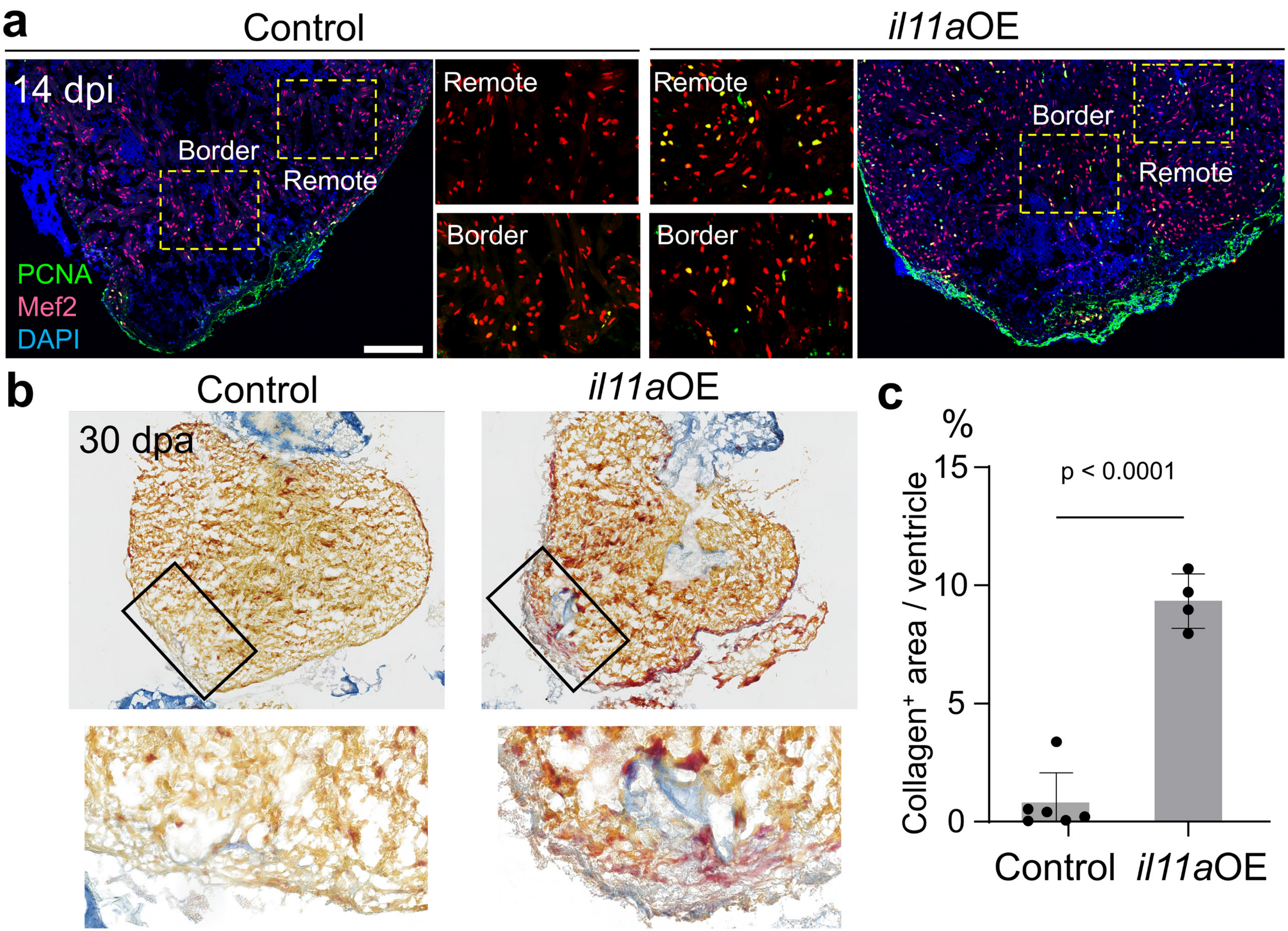
a
Control
il11aOE
14 dpi
Border
Remote
PCNA
Mef2
DAPI
Remote
Border
Remote
Border
Border
Remote
b
Control
il11aOE
30 dpa
c
%
Collagen+ area / ventricle
p < 0.0001
15
10
5
0
Control
il11aOE

**Supplementary Fig. 9. Dual roles of *il11a* in injured hearts.**

**(a)** CM proliferation in the regenerating hearts is enhanced by *il11a*OE. Representative image of 14 dpi heart sections stained with Mef2 (red) and PCNA (green) from control and *il11a*OE following 4-HT treatment. dpi, days post cryoinjury. Yellow dash boxes correspond to the region magnified in the border zone and remote area from control and *il11a*OE. **(b)** Representative AFOG staining images of control and *il11a*OE ventricles at 30 dpa. Bottom images correspond to higher magnifications of boxes. (**c**) Quantification of collagen$^+$ area in the ventricle of control and *il11a*OE at 30 dpa. n = 4 - 6. Scale bar, 100 μm in **a**.

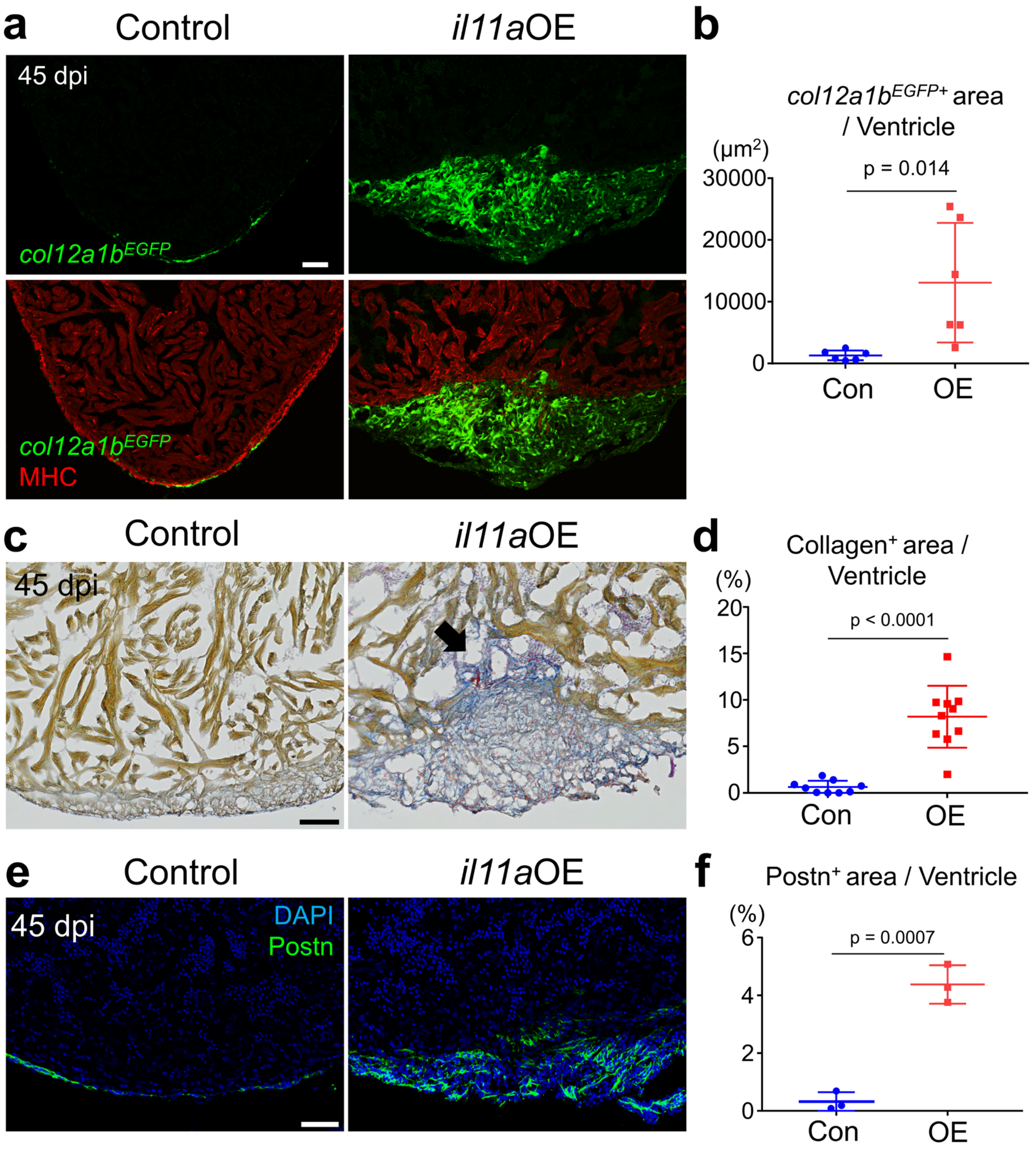
a
Control
il11aOE
45 dpi
col12a1bEGFP
col12a1bEGFP
MHC
b
col12a1bEGFP+ area / Ventricle
(µm2)
p = 0.014
30000
20000
10000
0
Con
OE
c
Control
il11aOE
45 dpi
d
Collagen+ area / Ventricle
(%)
p < 0.0001
20
15
10
5
0
Con
OE
e
Control
il11aOE
45 dpi
DAPI
Postn
f
Postn+ area / Ventricle
(%)
p = 0.0007
6
4
2
0
Con
OE

**Supplementary Figure 10. *il11a*OE leads to the persistence of EPC-derived fibroblasts and collagen deposition at the injury site after cryoinjury.**
(**a**) Representative cardiac section images of 45 dpi control and *il11a*OE expressing *col12a1b:EGFP*. Green and red indicate *col12a1b:EGFP* and MHC, respectively. (**b**) Quantification of $EGFP^+$ area at the injury site from Con and *il11a*OE. n = 6. (**c**) Representative AFOG staining images of control and *il11a*OE hearts at 60 dpi. (**d**) Quantification of $collagen^+$ area in the ventricle of Con and *il11a*OE at 60 dpi. The arrow indicates the presence of scar tissue near the injury site. n = 9 - 10. (**e**) Representative cardiac section images of 45 dpi control and *il11a*OE. Green represents $Postn^+$ fibroblasts at the injury site. (**f**) Quantification of $Postn^+$ area at the injury site from Con and *il11a*OE. n = 3. Scale bars, 50μm in **a, c,** and **e**

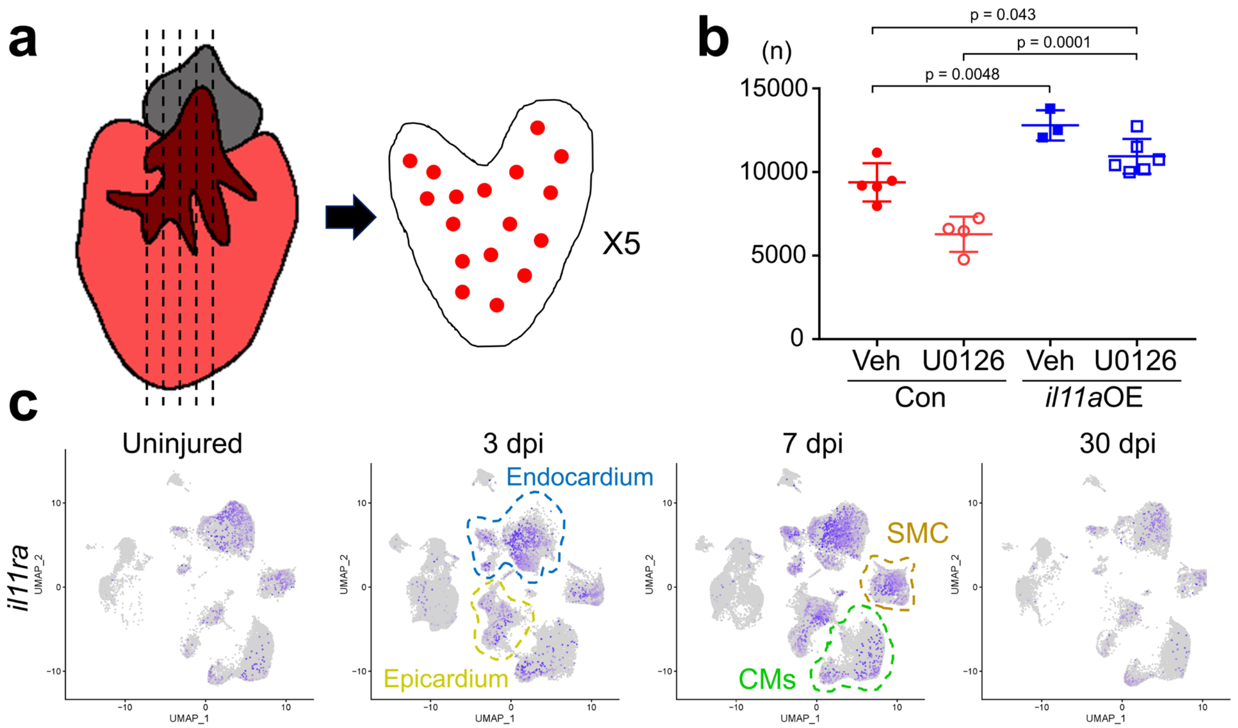


**Supplementary Fig. 11. Combinatorial treatment enhances cardiac regeneration in the zebrafish hearts.**
(**a**) Experimental design for counting CMs. 5 cardiac sections are selected (left) and Mef2$^+$ cells (red dots in cartoon) are quantified. (**b**) The total number of Mef2$^+$ cells in the entire ventricular region of 5 serial cardiac sections from veh or U0126-treated control and *il11a*OE hearts. n = 3 - 6. (**c**) Gene expression plots of *il11ra* displayed epicardial and endocardial expression at 3 days post injury (dpi), then expression in CMs and SMCs at 7 dpi.

**Table S1. Primer list**

| *il11a* overexpression line generation | |
|---|---|
| il11a ATG XmaI -f | cga cccggg atgaaattgctgggtgactcctcc |
| il11a stop NotI -r | aac gcggccgc ctatttccccacaattcgaatc |

| *il11a* knock-in reporter line generation | |
|---|---|
| il11a 5' HM HindIII -f | cgg aagctt acagactgctgtctcaggac |
| il11a 5' HM EcoRI -r | gcc gaattc caagtccttgttttaaaaggt |
| il11a 3' HM NotI -f | tcc gcggccgc gctctgttatatttgtttacatttagt |
| il11a 3' HM KpnI -r | ctt ggtacc tgagtgctggatgtgagcacaa |
| il11a start GG-2 sgRNA template | GCG TAATACGACTCACTATA GG atc aagtg ttact cgctc GTTTTAGA GCTAGAAAtagc |
| 3' universal primer | AAAGCACCGACTCGGTGCCACTTTTTCAAGTTGATAACGGACTAGCCTTATTTTA ACTTGCTATTTCTAGCTCTAAAAC |

| *In situ* hybridization primers | |
|---|---|
| socs3b ISH -f | CTTTCTCCTGGAAGGATGGAGCA |
| socs3b ISH -r | TCAGTGAATAGCAGACGTCCTG |
| jak1 ISH -f | ACTCTGCTCAACTATTCTGTGCA |
| jak1 ISH -r | TGTCAAGCATCTGCTGAAAGTT |
| stat3 qPCR -f | TGGGTCGAGAAGGACATCA |
| stat3 ISH -r | TTTGGCTCGGAGAGAGAAAG |
| angptl2a ISH -f | GGAGCCAGAGGCAGATTTCTACAA |
| angptl2a ISH -r | TGGAAAGTGTTGGGATTTGGTCGG |
| sulf1 ISH -f | CTATGGAAATCAAGCAGCTGGAGT |
| sulf1 ISH -r | CAGCTTTCAAAAAGGGCAAAATCC |

| | |
|---|---|
| thsd7aa ISH -f | AGTGTTACCTGACAGACTGGACGA |
| thsd7aa ISH -r | TTTCCCATCGGGACCAAAAGGTTG |

**Supplementary Data 1. Table of differentially expressed genes in control and 7 dpt *il11a*OE hearts**

**Supplementary Data 2. Table of differentially expressed genes in control and 7 mpt *il11a*OE hearts**
